# Impacts of dietary fat on multi tissue gene expression in the desert-adapted cactus mouse

**DOI:** 10.1101/2024.05.03.592397

**Authors:** Danielle M. Blumstein, Matthew D. MacManes

## Abstract

Understanding the relationship between dietary fat and physiological responses is crucial in species adapted to arid environments where water scarcity is common. In this study, we present a comprehensive exploration of gene expression across five tissues (kidney, liver, lung, gastrointestinal tract, and hypothalamus) and 19 phenotypic measurements, investigating the effects of dietary fat in the desert-adapted cactus mouse (*Peromyscus eremicus*). We show impacts on immune function, circadian gene regulation, and mitochondrial function for mice fed a lower-fat diet compared to mice fed a higher-fat diet. In arid environments with severe water scarcity, even subtle changes in organismal health and water balance can affect physical performance, potentially impacting survival and reproductive success. The study sheds light on the complex interplay between diet, physiological processes, and environmental adaptation, providing valuable insights into the multifaceted impacts of dietary choices on organismal well-being and adaptation strategies in arid habitats.

## Introduction

Harsh environments (e.g., high altitude, deep sea, deserts) represent natural experiments, given that organisms living in these extreme environments have evolved physiological, biochemical, and genomic mechanisms to persist (Blumstein and MacManes, 2023; Blumstein and MacManes, 2024; Kordonowy et al., 2017; Tigano et al., 2022; Wilsterman and Cheviron, 2021). One way for organisms survive under extreme conditions is to carefully manage their intake of food by making strategic decision based on environmental conditions which include food availability. Understanding how animals respond opportunistically to diet variation in extreme habitats can provide insight into the adaptive mechanisms they have evolved as well as the potential for other species to adapt to increasingly erratic climatic patterns. As the planet continues to undergo unprecedented climate change, increased desertification and a decrease in free water availability is predicted (IPCC, 2019; Mirzabaev et al., 2019). Several studies have demonstrated or predicted distributional changes in extant species in response to changing climate (Moritz et al., 2008; Parmesan, 2006), but fewer have examined the physiological (Bijlsma and Loeschcke, 2005; Blumstein and MacManes, 2023; Blumstein et al., 2024; Gabriel, 2005; Ramirez et al., 2022) and genomic (Blumstein and MacManes, 2024; Cheviron et al., 2014; Colella et al., 2021; MacManes, 2017; Tigano et al., 2020) mechanisms that may allow animals to adapt in place.

Animals consume three primary macromolecules—carbohydrates, fats, and proteins— which exhibit variation in water content (Mellanby, 1942). The oxidation of carbohydrates yields 0.60 g of metabolic water per gram, fats yield 1.07 g per gram, attributed to the higher oxygen demands of lipid metabolism, and proteins yield the least metabolic water (0.41 g) among macronutrients, requiring water loss for the excretion of nitrogenous waste (Davidson and Passmore, 1963; Mellanby, 1942; Schmidt-Nielsen and Adolph, 1964). Energy content also varies significantly, with carbohydrates and proteins providing 16.74 kJ/g, while fats offer more than double the energy at 37.66 kJ/g (Sánchez-Peña et al., 2017). Further, dietary composition is linked to metabolic water production, generated through the endogenous catabolism of carbohydrates, fats, or proteins, also displays substantial variation. In deserts, where water is scarce, both dietary composition and metabolic water production are crucial for survival.

In deserts, the stressors related to the lack of water are often further compounded by extreme temperatures which increases the rate at which water is lost via evaporation. If the lack of water and extreme temperatures don’t require more energy than an organism’s ability to absorb from the environment, the organism will maintain energy homeostasis (Romero, 2004). In arid environments such as deserts the interplay of both preexisting (dietary) water and internally generated water becomes crucial for survival. Food serves as the foundational resource for generating metabolic water but incurs costs during the breakdown of food leading to water loss through urination and fecal production (Schmidt-Nielsen, 1975). In desert ecosystems, rodents primarily subsist on vegetation, seeds, and insects, each exhibiting variations in fat, protein, and carbohydrate content (Orr et al., 2015; Wolf and del Rio, 2003). However, the specific composition of the diet is often unpredictable and highly diverse, especially for wildlife. Factors such as energy potential, nutritional content, and the capacity for water production (Schmidt-Nielsen and Adolph, 1964) are all contingent on the composition of the diet.

Faced with the natural variation in resource availability, opportunistic animals have the capacity to significantly impact their internal water balance and in turn, their survival. When comparing a standard diet to a low fat diet, Blumstein et al. (2024) described elevated rates of water loss during the warmer, drier light phase compared to the dark phase and variations in serum electrolyte levels suggestive of dehydration for cactus mice fed a low fat diet, suggesting dietary fat is significant in regulating water balance during long-term physiological experiments. Even minor variation in water balance can profoundly affect cognitive function (Riebl and Davy, 2013), physical performance, and ultimately, survival and reproductive success (Cunningham et al., 2021).

Metabolic flexibility is dependent on the capacity of reciprocal regulation of glucose use and fatty acid oxidation in the mitochondria (Muoio, 2014). When glucose consumption increases, fatty acid oxidation is suppressed, and vice versa (Hue and Taegtmeyer, 2009). During the postprandial phase following a carbohydrate-rich meal, glucose uptake, glycolysis, and pyruvate oxidation are favored due to elevated glucose and insulin levels (Muoio, 2014). Consequently, fatty acids are directed towards triglyceride synthesis and storage. Moreover, the availability of fat inhibits glycolysis, hindering glucose uptake and utilization.

During water deprivation, animals have been observed reduce solid food intake, a phenomenon known as dehydration-associated anorexia, which minimizes water needed for digestion and allows for water reabsorption from the kidneys and gastrointestinal tract (Armstrong et al., 1980; Blumstein and MacManes, 2024; Hamilton and Flaherty, 1973; Rowland, 2007; Salter and Watts, 2003; Schoorlemmer and Evered, 2002; Watts and Boyle, 2010). Glucose levels are maintained during limited food intake and water deprivation due to enhanced glycogenolysis and gluconeogenesis (Blumstein and MacManes, 2024; Salter and Watts, 2003; Schoorlemmer and Evered, 2002; Watts and Boyle, 2010). The inhibition of glycolysis impedes glucose metabolism (Spriet, 2014). This regulatory mechanism creates a positive feedback loop that promotes fatty acid oxidation during nutrient scarcity, preserving glucose for essential biosynthetic processes and brain metabolism.

Food intake and composition also servers a twofold function, acting as both a source of energy and a regulator of the gastrointestinal tract (Henderson et al., 2015; Keeney and Finlay, 2011; Wu et al., 2011). Nutrients, obtained from the host, play a crucial role in shaping the composition of host-associated microbial communities (McFall-Ngai, 2007). Despite the mutually beneficial relationship between intestinal microbes and the host, the proximity of an abundant bacterial community to intestinal tissues poses significant health risks. Thus, the gut microbiota plays a crucial role in diet processing but simultaneously, dietary changes can influence the composition of the gut microbial community (Sonnenburg et al., 2016).

To fully understand the mechanisms that control organismal health, responses must be studied in parallel and at multiple levels. We extend previously characterized metabolic patterns for males and females cactus mice fed an experimental diet low in fat to those on a standard laboratory diet (Blumstein et al., 2024) with a thorough examination of gene expression patterns in five tissues (kidney, liver, lung, gastrointestinal tract, and hypothalamus). We describe the whole-organism physiological and genomic response to variations in dietary fat in desert adapted opportunistic animals. Overall, the gastrointestinal tract experienced the largest number of changes in gene expression, followed by lung, the hypothalamus, kidney, and liver. The WGNCA analysis of the kidney, liver, and lung revealed significant modules related to circadian rhythm while the hypothalamus, kidney, liver, and lung all had significant differentially expressed genes related to mitochondrial gene expression. Additionally, we highlight the significance of dietary fat in immune response. These findings suggest that a low-fat (LF) diet may limit the survival of the cactus mouse in the event of restricted access to water, a common challenge in arid habits.

## Methods

### Animal Care and RNA Extraction

Captive-bred, sexually mature, non-reproductive healthy male and female *P. eremicus* raised in an environmental chamber replicating the climate of the Sonoran Desert were used in this study. We adhered to animal care procedures established by the American Society of Mammologists and sanctioned by the University of New Hampshire Institutional Animal Care and Use Committee (protocol number 210604). The mice (n = 28 males, n = 28 females), provided water ad libitum, were randomly assigned to one of two diet treatment groups, a standard diet group (SD - LabDiet® 5015*, 26.101% fat, 19.752% protein, 54.148% carbohydrates, energy 15.02 kJ/g, food quotient [FQ, the theoretical RQ produced by the diet based on macronutrient composition, [Westerterp, 1993] 0.89 or a low-fat diet group (LFD - Modified LabDiet® 5015 [5G0Z], 6.6% fat, 22.8% protein, 70.6% carbohydrates, energy 14.31 kJ/g, FQ = 0.92)

Prior to the beginning of the experiment, mice were fed the assigned diet for four weeks. We weighed the mice then transferred them to an experimental cage 24 hours prior to the beginning of metabolic measurements, then initiated undisturbed data collection for 72 hours. Throughout the study, pull flow-through respirometry system (Lighton, 2018) was used to assess metabolic phenotypes, computing CO_2_ production rates, O_2_ consumption, and water loss using established equations. At the conclusion of the experiment (day 3), weight was again recorded, the mice were euthanized, and trunk blood was collected and electrolytes data were collected using an Abaxis i-STAT® Alinity machine and CHEM8+ cartridges (Abbott Park, IL, USA, Abbott Point of Care Inc.) we measured the concentration of sodium (Na), potassium (K), creatinine (Cr), blood urea nitrogen (BUN), hematocrit (HCT), glucose (Glu), and ionized calcium (iCa), which are expected to vary in response to hydration status and renal function. Finally, using sodium, glucose, and BUN, we calculated osmolality using the formula in Rasouli (2016).

Following similar methods to Blumstein and MacManes (2024), the lung, liver, kidney, a section of the GI tract, and hypothalamus were collected, preserved, and RNA was extracted from anatomically similar regions of the tissue using a standardized Trizol protocol. RNAseq libraries were prepped for sequencing using a poly-A tail purification, barcoded using Illumina primers, and dual-barcoded using a New England Biolabs Ultra II Directional kit (NEB #E7765). Individually labelled samples were pooled and sequenced using a paired end protocol and a read length of 150bp, across two lanes of a Novaseq 6000 at the University of New Hampshire Hubbard Center for Genome Studies.

### Computational Analysis

The processed reads were aligned to the *P. eremicus* genome version 2.0.1 from the DNA Zoo Consortium (dnazoo.org) using STAR version 2.7.10b (Dobin et al., 2013), and aligned reads were counted using HTSEQ-COUNT version 2.0.2 (Anders et al., 2015). Count files were imported into R v 4.0.3 (R Core Team, 2020). Counts were merged into a gene-level count by combining all counts that mapped to the same gene. Low expression genes were removed if the gene had 10 or less counts in 8 or more individuals across the experiment.

Differential gene expression analysis was done using DESEQ2 (Love et al., 2014) on the dataset as a whole as well as for each tissue independently, with three different models that tested for the effects of sex, diet, and tissue type and for each tissue independently with two models, testing the effect of and identifying genes specific to sex and diet with a Wald test. We used a Benjamini and Hochberg (1995) correction for multiple comparisons and used an alpha value of .05 for significance. This was followed by weighted gene correlation network analysis (WGCNA) (Langfelder and Horvath, 2008) for each tissue independently and included all the physiological phenotypes collected (mean EE, water loss, RQ, weight, sex, diet, and the panel of electrolytes). Canonical Correlation Analysis (CCA) in the R package vegan (Oksanen, 2010) was performed to explore gene expression correlations across tissues, diet, sex, and metabolic variables. ANOVA was used to identify what response variables were significant for graphing on the triplot, allowed us to visualize their correlative nature; vectors pointing in the same direction are positively correlated, while vectors pointing in opposite direction are negatively correlated. To identify genes of interest, we selected genes that graphed two standard deviations away from the origin for CCA1 and CCA2. Lastly, gene ontology analyses were conducted using the R package gprofiler (Kolberg et al., 2023) and identified the topmost 20 significant GO terms based on g:SCS corrected p-values (Reimand et al., 2007) to elucidate significant gene functions and pathways associated for each up and downregulated list for each tissue and for each significant module from the individual WGCNA analysis. The full analytical code and data are available on the GitHub repository (https://github.com/DaniBlumstein/diet_rnaseq). Finally, we compiled a list of genes with high confidence from the findings of two of the independent analyses: WGCNA and genes located two standard deviations from the origin in the CCA.

## Results

All mice underwent routine animal care, including health evaluations conducted by licensed veterinary professionals, prior to the experiment. No health concerns were identified during these assessments and none of the animals were removed prematurely. All mice remained active for the duration of the experiment.

### Genomic Data

On average, we generated 13.76 million reads per sample, with a standard deviation of 35.08 million reads. On average, 78.15% of reads per sample were uniquely mapped, with a standard deviation of 0.03%. Detailed information regarding the number of reads and per-sample mapping rates can be found in Supplemental File 1. Raw read files are archived at NCBI SRA BioProject: PRJNA1048512, and the gene expression count data along with the analytical code are accessible in the GitHub repository (https://github.com/DaniBlumstein/diet_rnaseq).

### Electrolytes and Physiological Phenotypes

The same mice used to generate the electrolyte and physiology data more fully described in (Blumstein et al., 2024) are also used in the study described herein for RNAseq analysis. There were no significant differences in weight between males and females (mean of 22.2 g [range: 15.40 - 29.10 g] versus 21.2 g [range: 16.91 - 29.34 g], respectively; Table 1), nor between diet groups. When comparing males and females separately, the following electrolytes were significantly different for the SD vs the LFD treatments: sodium p = 0.22 and p = 0.01 respectively; potassium p = 0.02 and p = 0.01; BUN p = 0.03 and 0.71; hematocrit p = 0.04 and 0.02; ionized calcium p = 0.001 and 0.001; osmolality p = 0.01 and 0.05 (Table 1). When comparing males and females on the same diet, the following electrolytes were significantly different: sodium (p = 0.02) and osmolality (p = 0.03) differed significantly for mice on the SD (Table 1). No electrolyte measurements were differed between males and females on the LFD (Table 1).

Using data collected in in (Blumstein et al., 2024), we calculated the means for EE, WLR, and RQ for the same 28 adult females and 28 adult males (n=14 of each treatment, total n=56) for each mouse from the last hour of data collected . Data located at: <u>https://github.com/DaniBlumstein/Diet_paper</u>. Within sex, EE significantly differed between diets for males (p = 0.012) but not females (p = 0.167), WLR was significantly different between diets for males (p = 0.001) and females (p = 2.641e-05). RQ was not significantly different.

**Table 1.** Summary statistics for weight (g) and serum electrolyte measurements (Na = Sodium (mmol/L), K = Potassium (mmol/L), Cr = Creatinine (μmol/L), BUN = Blood Urea Nitrogen (mmol/L), Hct = Hematocrit (% PCV), iCa = Ionized Calcium (mmol/L), and osmolality (mmol/L) for female and male *Peromyscus eremicus* for standard and low-fat diet groups. Bold values represent significant differences within sex between diet while italicized values represent significant differences between sex within diet.

|  | Standard diet |  |  | Low-fat diet |  |  |
| --- | --- | --- | --- | --- | --- | --- |
|  | Female | Male | All | Female | Male | All |
| Weight (g) | 22.4+/-2.73 | 22.24+/-2.14 | 21.13+/-2.63 | 20.02+/-2.68 | 22.29+/-4.05 | 22.35+/-3.38 |
| Na (mmol/L) | **137.5+/-3.58** | **140.71+/-2.89** | 139.23+/-3.56 | ***143.6+/-4.7*** | ***142.5+/-3.63*** | 143.05+/-4.12 |
| K (mmol/L) | **8.02+/-1.35** | **7.84+/-0.96** | 7.92+/-1.14 | **5.98+/-1.49** | **6.37+/-1.59** | 6.18+/-1.51 |
| Cr (μmol/L) | 0.22+/-0.04 | 0.2+/-0.03 | 0.21+/-0.03 | 0.2+/-0 | 0.2+/-0 | 0.2+/-0 |
| BUN (mmol/L) | 36.25+/-5.33 | **33.21+/-4.08** | 34.62+/-4.85 | 38.3+/-5.5 | **37+/-3.53** | 37.65+/-4.55 |
| Hct (% PCV) | **26.08+/-9.58** | **32.5+/-2.79** | 29.54+/-7.42 | **34.4+/-3.27** | **36.2+/-5.57** | 35.3+/-4.54 |
| iCa (mmol/L) | **1.1+/-0.1** | **1.16+/-0.07** | 1.13+/-0.09 | **1.29+/-0.07** | **1.27+/-0.09** | 1.28+/-0.08 |
| Osmolality (mmol/L) | **291.15+/-7.34** | **297.54+/-5.82** | 295.04+/-7.1 | **307.83+/-10.3** | **304.18+/-9.22** | 306.01+/-9.7 |

### Differential Gene Expression

Following the methods in Blumstein and MacManes (2024), we cross-referenced our gene IDs with *Homo sapiens* gene IDs and filtered low expression genes, leaving 17012 genes in the dataset as a whole. Patterns of gene expression data are largely driven by tissue type (PC1: 44% variance and PC2: 23% variance, Supplemental Figure 1).

All downstream analysis, with the exception of CCA (see below) were done one tissue at a time by filtering the count and sample data files by the tissue of interest for that analysis. This left 12935 genes in the lung, 10820 genes in the liver, 12249 genes in the GI, 13037 genes in the hypothalamus, and 11950 genes in the kidney. Each tissue had a unique number of differentially expressed genes (p < 0.05) between diet treatment and sex (Figure 1, Table 2).

**Figure 1.**
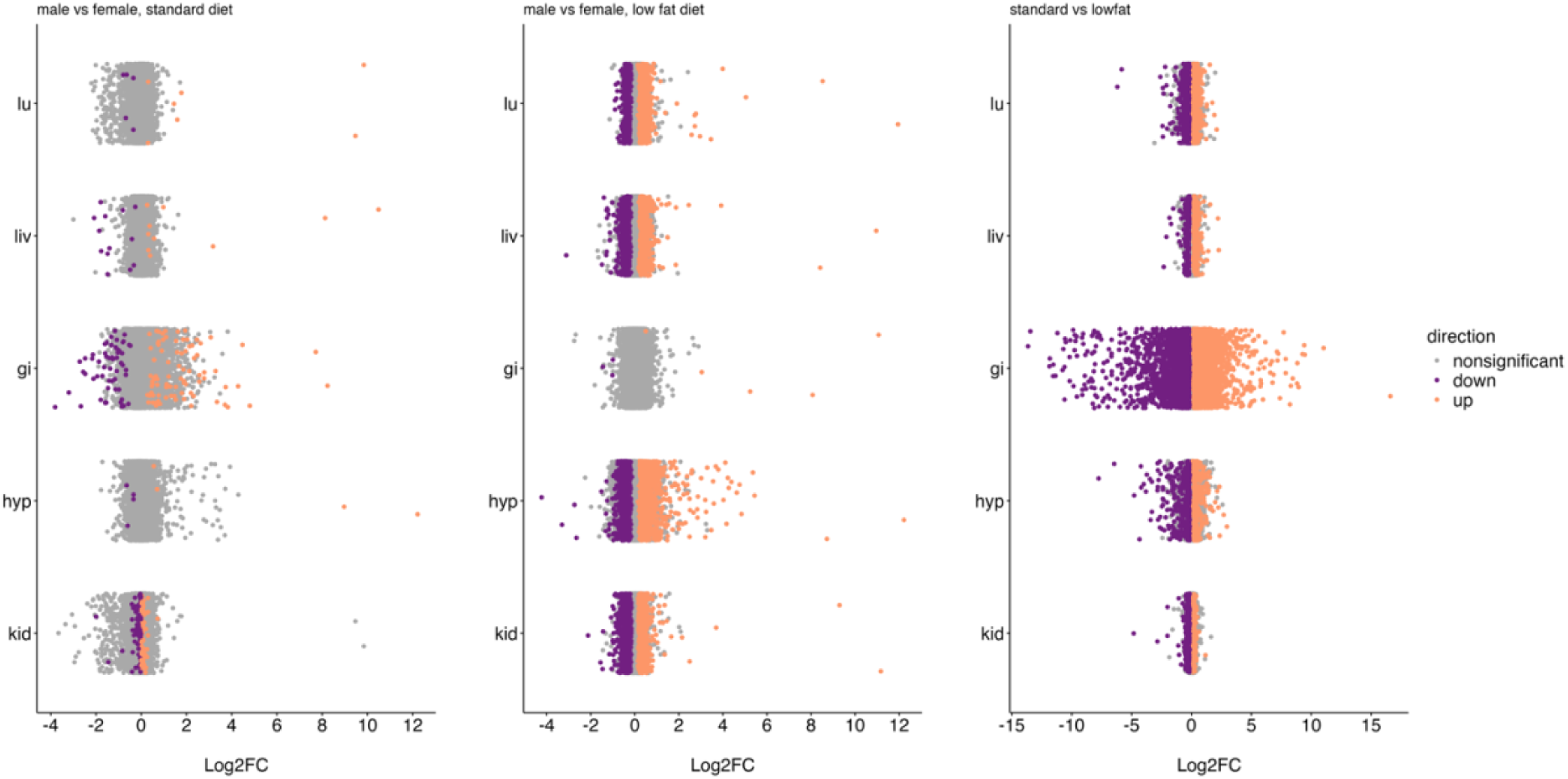
Log fold change of all genes across the lung (lu), liver (liv), gastrointestinal tract (gi), hypothalamus (hyp), and kidney (kid) of *Peromyscus eremicus* comparing males versus females on the standard diet, males versus females on the low-fat diet, and all mice the standard diet to all mice on the low-fat diet. Purple (downregulated) and orange (upregulated) colored dots indicate p<0.05, whereas grey dots indicate p>=0.05.

**Table 2.** The number of differentially expressed (DE) genes in the lung, liver, gastrointestinal tract, hypothalamus, and kidney of *Peromyscus eremicus* for males versus females and the standard diet versus low-fat diet (adjusted p < 0.05).

|  | DE between sexes (SD)<br>males vs females |  | DE between sexes (LF)<br>males vs females |  | DE between diet treatments<br>standard vs low-fat |  |
| --- | --- | --- | --- | --- | --- | --- |
|  | upregulated | downregulated | upregulated | downregulated | upregulated | downregulated |
| lung | 7 | 5 | 742 | 452 | 1074 | 1123 |
| liver | 10 | 13 | 663 | 446 | 246 | 200 |
| GI | 84 | 57 | 5 | 3 | 3652 | 3095 |
| hypothalamus | 4 | 4 | 978 | 744 | 277 | 503 |
| kidney | 87 | 45 | 1017 | 815 | 178 | 249 |

### Weighted Gene Correlation Network Analysis

To better link within-tissue patterns of gene expression with treatment, condition, or collected phenotypic data, we conducted an analysis in WGCNA. Briefly, WGCNA is a network-based statistical approach that identifies clusters of genes with highly correlated expression profiles (modules), or genes that could function together in a pathway (Langfelder and Horvath, 2008). For the lung, we identified 30 modules, of which 23 were significant, with each module containing 20-8292 genes. Notably, nine modules were significant for three or more phenotypes. In the liver, 40 modules were identified, 26 of which were significant and contained 20-14458 genes per module. Seven of these modules were significant for three or more phenotypes. In the gastrointestinal tract, 32 modules were identified, and 25 were significant, each with 21-7599 genes. Fifteen modules were significant for three or more phenotypes. In the hypothalamus, we found 21 modules, of which 18 were significant, and they contained 23-9472 genes per module. Ten modules were significant for three or more phenotypes. Finally, in the kidney, we identified 37 modules, 25 of which were significant and contained 21-13762 genes per module. Seven of these modules were significant for three or more phenotypes. Several phenotypes (Na, Bun, AnGap, K, TCO2, iCa, diet treatment, and EE) had at least one significant module in all five tissues. There was at least one significant module for sex in the hypothalamus, liver, kidney, and lung. At least one significant module for Hb in the hypothalamus, liver, GI, and lung. At least one significant module for RQ in the hypothalamus, liver, GI, and lung. At least one significant module for Cl in the hypothalamus, kidney, GI, and lung. At least one significant module for Hct in the liver, GI, and lung. At least one significant module for glucose in the kidney, GI, and lung. At least one significant module for WLR in the hypothalamus, GI, and lung. At least one significant module for weight in the liver and kidney and there were no significant modules for creatin in any of the tissues.

### Gene Ontology

To enable functional interpretation, we filtered to the top 20 Gene Ontology (GO) terms, based on p values, for the upregulated and downregulated significantly differentially expressed genes for each tissue. We observed a total of 343 unique GO terms. Of these, 78 GO terms are upregulated genes (e.g. GI: wound healing [GO:0042060], hypothalamus: neurogenesis [GO:0022008]) and 76 are downregulated genes in the diet comparison (e.g. GI: fatty acid catabolic process [GO:0009062] and carbohydrate catabolic process [GO:0016052], hypothalamus: mitochondrion organization [GO:0007005] and mitochondrial electron transport, NADH to ubiquinone [GO:0006120], kidney: fatty acid metabolic process [GO:0006631] and mitochondrial envelope [GO:0005740], liver: mitochondrion [GO:0005739]). When comparing males versus females we analyzed the two diets separately and found 30 GO terms upregulated for the SD (e.g. GI: T cell differentiation [GO:0030217], activation of immune response [GO:0002253], and response to stimulus [GO:0050896]), 35 terms downregulated for the SD (e.g. liver: rhythmic process [GO:0048511], lung: rhythmic process [GO:0048511]), 66 terms upregulated for the LF (hypothalamus: carbohydrate derivative metabolic process [GO:1901135], mitochondrial large ribosomal subunit [GO:0005762], liver: import into the mitochondrion [GO:0170036], lung: mitochondrial translation [GO:0032543]), and 58 downregulated for the LF (e.g. hypothalamus: autophagosome assembly [GO:0000045], liver circulatory system development [GO:0072359] and regulation of response to stimulus [GO:0048583], lung: circulatory system development [GO:0072359] and regulation of response to stimulus [GO:0048583], (Figure 2).

GO analysis of the genes contained in significant WGCNA modules identified a total of 455 unique GO terms. Among these terms, 61 GO terms were found to be present in two tissues, 16 in three tissues (mitochondrial matrix [GO:0005759], circadian rhythm [GO:0007623], photoperiodism [GO:0009648], entrainment of circadian clock [GO:0009649], circadian regulation of gene expression [GO:0032922], regulation of circadian rhythm [GO:0042752], entrainment of circadian clock by photoperiod [GO:0043153], rhythmic process [GO:0048511], cytoplasmic vesicle [GO:0031410], isoprenoid metabolic process [GO:0006720], secretory granule [GO:0030141], cell projection organization [GO:0030030], plasma membrane [GO:0005886], cell periphery [GO:0071944], catalytic activity [GO:0003824], small molecule metabolic process [GO:0044281]), one in four tissues (cellular lipid metabolic process [GO:0044255]), and one in five tissues (cytoplasm [GO:0005737]) (Figure 2). Specifically, in the lung, 143 unique GO terms were identified, with 136 terms in one module, six terms in two modules (secretory vesicle [GO:0099503], lytic vacuole [GO:0000323], cell activation [GO:0001775], immune system process [GO:0002376], lysosome [GO:0005764], vesicle [GO:0031982]), and one term in four modules (immune response [GO:0006955]) (Figure 2). In the liver, 82 unique GO terms were found, with 78 terms in one module, and four terms in two modules (catalytic activity [GO:0003824], small molecule metabolic process [GO:0044281], oxidoreductase activity [GO:0016491], purine-containing compound metabolic process [GO:0072521]) (Figure 2). For the GI, 119 unique GO terms were identified, with 115 terms in one module and four terms in two modules (cytoplasm [GO:0005737], immune response [GO:0006955], regulation of signal transduction [GO:0009966], cell periphery GO:0071944) (Figure 2). In the hypothalamus, 89 unique GO terms were found, with 81 terms in one module, four terms in two modules (plasma membrane [GO:0005886], vesicle-mediated transport [GO:0016192], external encapsulating structure [GO:0030312], intracellular protein-containing complex [GO:0140535]), and four terms in three modules (cell periphery [GO:0071944], extracellular organelle [GO:0043230], extracellular membrane-bounded organelle [GO:0065010], extracellular vesicle [GO:1903561]) (Figure 2). Lastly, in the kidney, 118 unique GO terms were identified, with 111 terms in one module and seven terms in two modules (cellular lipid metabolic process [GO:0044255], extracellular exosome [GO:0070062], mitochondrion [GO:0005739], negative regulation of nucleobase-containing compound metabolic process [GO:0045934], regulation of cellular process [GO:0050794], protein-DNA complex organization [GO:0071824], plasma membrane region [GO:0098590]) (Figure 2).

**Figure 2.**
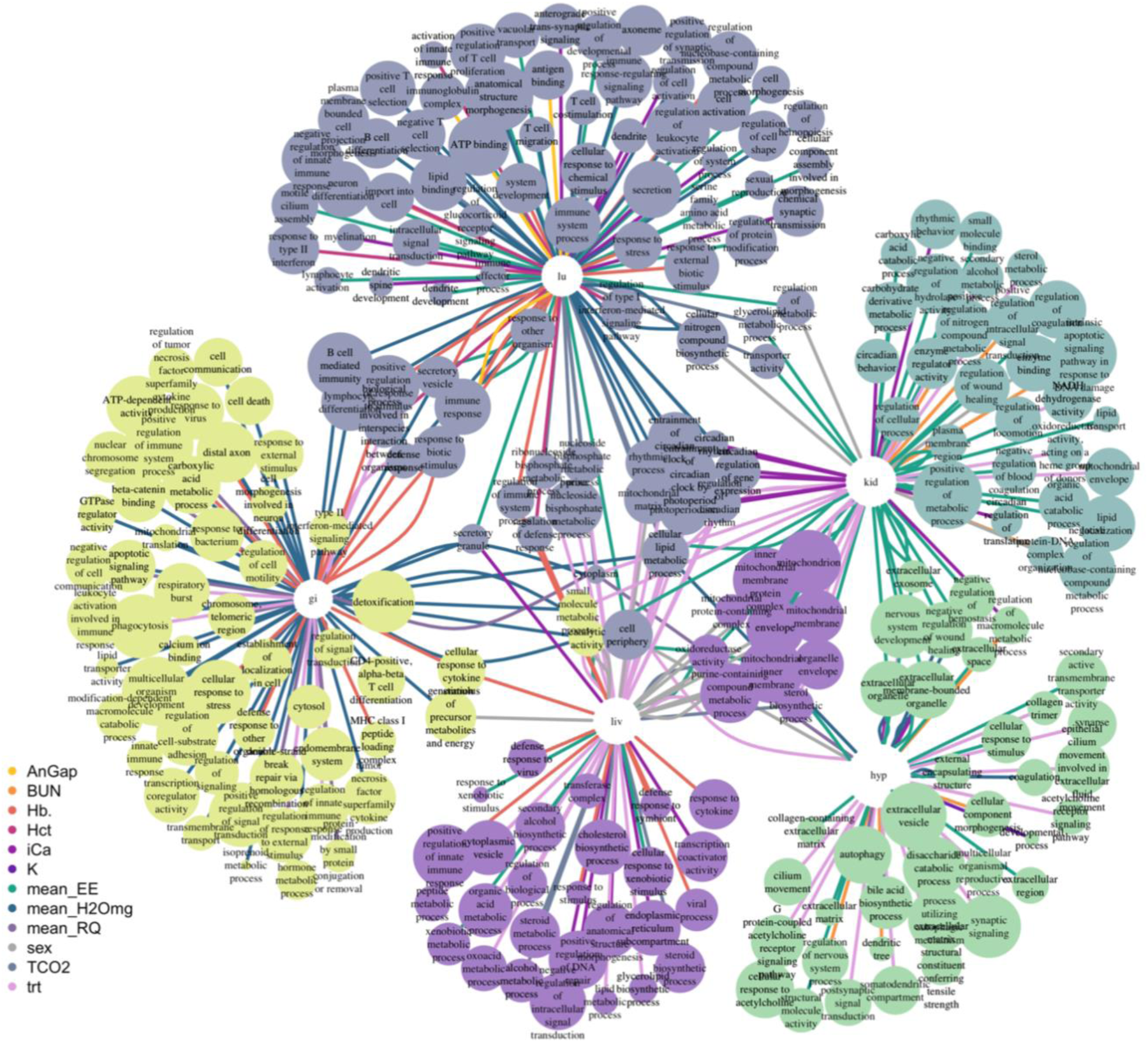
Visualization of gene ontology (GO) terms to show common WGCNA modules within and between the lung (lu), liver (liv), gastrointestinal tract (gi), hypothalamus (hyp), and kidney (kid) of *Peromyscus eremicus*. Visualized are selections of the top 20 significant GO terms for each phenotype module combination. The number of genes in the GO term are indicated by size of the node and the color of the edges represent the phenotype for that module. Clusters are placed according to the Kamada-Kawai algorithm (Kamada and Kawai, 1989) and are largely driven by tissue followed by shared GO terms between tissues.

### Canonical Correlation Analysis

We examined the relationship between gene expression and water access, physiological variables (EE, RQ, WLR, and weight), tissue type, and sex using CCA (Oksanen, 2010). CCA explained a large amount of variance (CCA1: 38.07% and CCA2: 20.09%, Figure 3). Additionally, 1296 genes were identified as being two standard deviations from the origin. Overall, the model was significant (p=0.001, Table 3) meaning that there is an association among our variables. Weight was significant (F = 2.44, p= 0.046, Table 3) with the vector pointing at the kidney, RQ was significant (F = 4.02, p = 0.004, Table 3) with the vector pointing at the hypothalamus, and EE was significant (F = 3.57, p = 0.004, Table 3) with the vector pointing at the GI. This suggests that gene expression in the hypothalamus, kidney, and the GI effects RQ, weight, and EE differently.

**Table 3.** ANOVA results from the Canonical Correspondence Analysis showing that diet (trt), weight, RQ (mean_RQ), EE (mean_EE), sex, tissue, and the CCA model are all significant. Formula: gene expression ∼ diet + weight + RQ + EE + WLR + sex + tissue).

|  | Df | ChiSquare | F | Pr(>F) |
| --- | --- | --- | --- | --- |
| trt | 1 | 0.00033 | 12.78 | 0.001 |
| weight | 1 | 0.00006 | 2.44 | 0.046 |
| mean_RQ | 1 | 0.00010 | 4.02 | 0.004 |
| mean_EE | 1 | 0.00009 | 3.57 | 0.004 |
| mean_H2Omg | 1 | 0.00005 | 1.83 | 0.116 |
| sex | 1 | 0.00012 | 4.82 | 0.001 |
| tissue | 4 | 0.03313 | 322.99 | 0.001 |
| Model | 10 | 0.03389 | 132.14 | 0.001 |
| Residual | 178 | 0.00457 |  |  |

**Figure 3.**
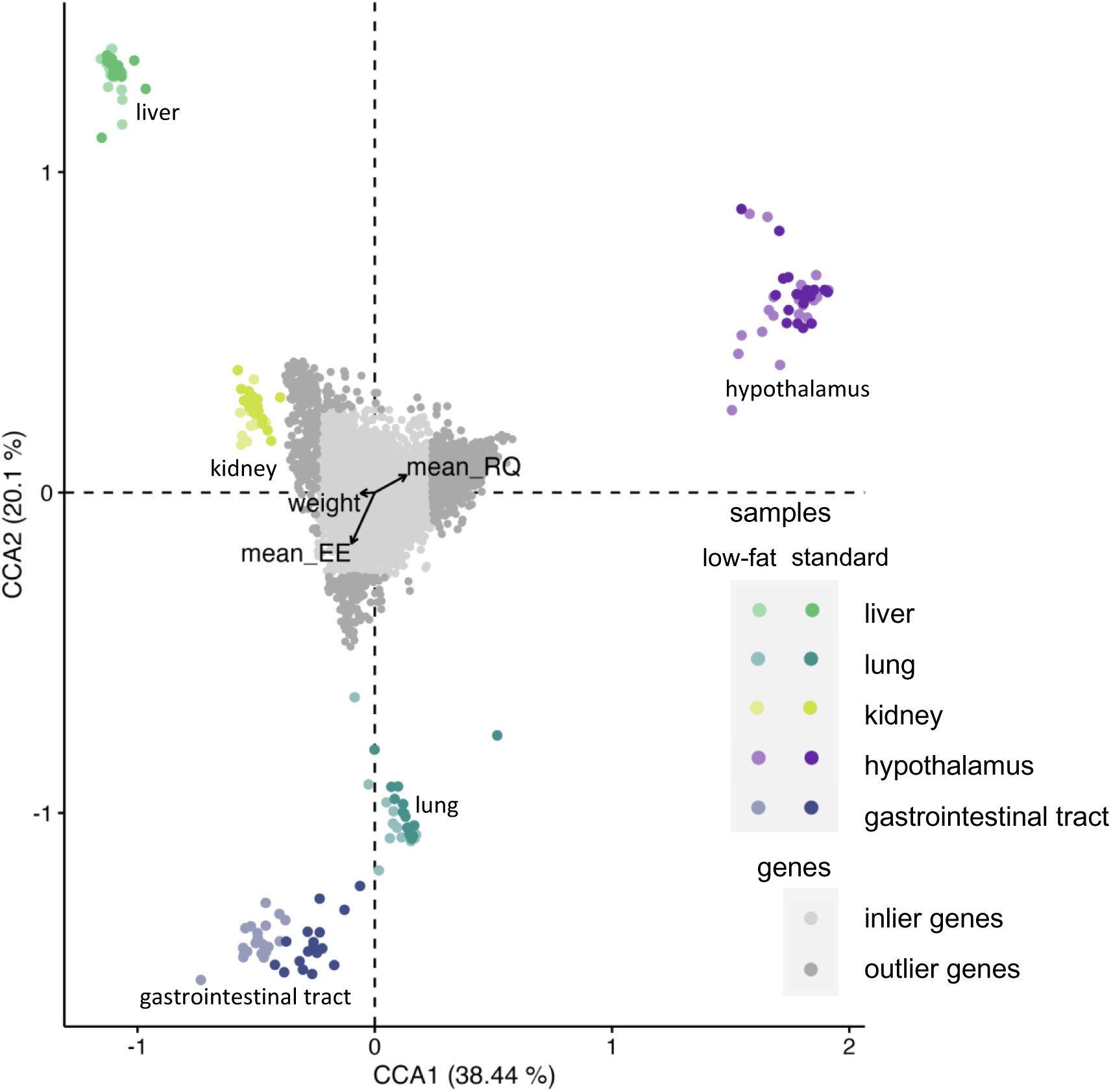
Canonical correspondence analysis (CCA) indicates correlations between normalized differentially expressed genes and physiological measurements for *Peromyscus eremicus* fed a standard diet and a low-fat diet. The distribution of tissue samples in Euclidian space as a function of their gene expression values is shown (points colored by tissue type and water treatment). Outlier genes (defined as two standard deviations or more from the mean) are colored dark grey. Inlier genes (defined as less than two standard deviations from the mean) are colored light grey. CCA reveals a significant relationship between weight (F = 2.44, p = 0.046), RQ (F = 4.02, p = 0.004), and EE (F = 3.57, p = 0.004).

### Consensus gene set

The overlap between CCA outliers, DE genes, and genes contained within significantly WGCNA module was examined for each tissue independently. For each gene set, we identified the function of each by performed independent GO and KEGG analyses (Supplemental Table 2). In the lung, there were 76 overlapping genes out of 12667 with functions related to apoptosis, immune, inflammation, angiogenesis, glutamate, membrane component, nervous system development, mitochondrial, and calcium (see Supplemental Table 2 for the full list). In the liver, 34 genes overlapped out of 10244 related to gluconeogenesis, immune system, insulin, etc. (Supplemental Table 2). In the kidney, 17 genes overlapped out of 11908 related to sodium, angiogenesis, apoptosis, gluconeogenesis, etc. (Supplemental Table 2). In the hypothalamus, 46 genes overlapped out of 11025 related to calcium, digestion and absorption, glutamate, immune system, mitochondrial, nervous system development, etc. (Supplemental Table 2). In the GI, 442 genes overlapped out of 12200 related to cholesterol, digestion and absorption, glutamate, inflammation, and insulin (Supplemental Table 2).

## Discussion

Currently, there is limited research on the effects of low-fat diet as studies have historically focused on the consequences of high fat diets (Abdel-Rahman, 2010; Hino et al., 2017; Kohsaka et al., 2007; Leone et al., 2015; Song et al., 2004). Previous work has explored diet composition and variation (Bachmanov et al., 2002; Beale et al., 2018; De la Iglesia et al., 2016; Hochman and P. Kotler, 2006; Hope and Parmenter, 2007; Nagy, 1994). Here we expand these to characterize the physiological and genomic mechanisms in response to a low-fat diet. This is of particular interest as diet directly affects both energy and water homeostasis (Schmidt-Nielsen and Adolph, 1964). Consumers respond to diets with multiple organ systems and physiological processes, a difficult signal to identify as gene expression highlights differences between tissues rather than processes coordinated by one or more tissue. It has been demonstrated that distributional changes of plants and consumers, in response to a changing climate, is occurring (Moritz et al., 2008; Parmesan, 2006), resulting in diet shits and unexpected organismal responses.

We investigated the impact of dietary fat in a desert-adapted rodent using a multi-tissue gene expression dataset with whole-organism physiological response to explore the interplay between physiological and genomic responses in harsh habitats. We observed that mice fed a lower-fat diet had 1) differences in expression levels in immune system genes, 2) gene expression changes in circadian regulating genes, and 3) expression changes related to mitochondrial function. In regions characterized by severe water scarcity, even subtle deviations of overall organismal immune function and water balance may significantly affect physical performance and consequently survival and reproductive success. Our findings contribute to a better understanding of the organismal significance of food availability and dietary preferences and suggest that survival might be less dependent on a high fat diet when water is accessible.

### Diet and dehydration

Blumstein et al. (2024) observed that mice on a diet lower in fat had a pronounced increase in water loss compared to those on a higher fat diet. While animals have *ad lib* access to water in this experiment, the differences in water loss may be important in arid environments where water scarcity is extreme. Even subtle differences in water balance could profoundly impact cognitive function, physical performance, and, consequently, survival and reproductive success (Cunningham et al., 2021; Riebl and Davy, 2013). Key electrolytes sensitive to hydration status, such as serum Na, osmolality, and Hct, are elevated in animals fed the LF diet indicating a difference in water balance. However, synthetic markers of pathological renal impairment, BUN, and Cr, did not significantly differ between treatments (further discussed in Blumstein et al., 2024). In the study described here, we found several significant genes identified in our consensus gene sets related to the management of Na (*SLC9A5* [lung], *TNR* [hypothalamus], *SLC38A4* and *SLC17A1* [kidney], Supplemental Table 2) and osmolality (*ADCYAP1R1* and *PKLR* [hypothalamus], *ITLN1* [lung], *SLC2A9* [liver], *GJB1* and *PPP1R3G* [kidney], Supplemental Table 2), suggesting that varied fat content results in a solute concentration response. However, genetic markers of renal impairment (see MacManes 2017) were not identified in the consensus gene set (Supplemental Table 2).

During dehydration, the renin-angiotensin-aldosterone system (RAAS) and vasopressin are upregulated for solute management (Aisenbrey et al., 1981; Bouby and Fernandes, 2003; Greenleaf, 1992; Roberts et al., 2011). Water deprived cactus mice have a consistent and robust whole-body RAAS response for water balance regulation (Blumstein and MacManes, 2024). In the current experiment, the primary genes responsible for this response *AGT* (angiotensinogen), *ACE2* (angiotensin-converting enzyme), *REN1* (Renin), *CMA1* (converts angiotensin I to angiotensin II), and *AGTR1* (Angiotensin II Receptor), were either not found in our datasets (*CMA1* and *REN1)* or were downregulated in LF mice (*AGT* and *AGTR1* [GI]), which is contrary to our prediction. Only *ACE2* and *AGTR1* [lung] were found to be upregulated in LF mice. Additionally, the vasopressin pathway (hsa04962, Kanehisa et al., 2023) is only partially differentially expressed, with *RAB5C*, *CREB3L2,* and *AQP3* significant upregulated while *AVPI1, RAB11B,* and *VAMP2* is significantly downregulated in LF mice in the GI. *DYNLL2* is significantly upregulated and *VAMP2* is significantly downregulated in LF mice in the lung. Lastly, *AVP* is significantly upregulated in LF mice in the hypothalamus. These responses are very different to those seen in Blumstein and MacManes (2024) where the RAAS pathway was found to be activated, and suggests that mice were not dehydrated to the levels that would stimulate a systematic and whole-organismal response.

The management of the consequences of altered fluid balance during dehydration involves processes like angiogenesis or vascular remodeling that maintain perfusion, addressing challenges from reduced fluid availability and potential hemoconcentration (discussed in Blumstein and MacManes, 2024). In the study described here, we found two upregulated genes identified in our consensus gene sets, *ANG* (lung), *ADTRP* (liver and kidney) involved in angiogenesis (Supplemental Table 2). However, a WGCNA module in the kidney grouping into the GO term blood vessel development [GO:0001568] were downregulated. Furthermore, genes related to coagulation factors during dehydration that mitigate risks associated with increased blood viscosity and clot formation were downregulated. We found one downregulated module in the hypothalamus related WLR to coagulation (coagulation [GO:0050817], and blood coagulation, fibrin clot formation [GO:0072378]) as well as two downregulated genes identified in our consensus gene sets, *F5* and *C8G* (GI) (Supplemental Table 2) related to coagulation. However, we also found three genes upregulated in the GI related to coagulation (*CFI*, *PLG*, and *F13B* Supplemental Table 2). This suggests that the repones to dehydration on a LF diet is not a global one, unlike what is discussed in Blumstein and MacManes (2024) because mice have water *ad libitum* and are not dehydrated.

Lastly, Blumstein and MacManes (2023; 2024) hypothesized water deprived mice reduce the amount of solid food intake in an response known as dehydration associated anorexia. This response reduces the amount of water required for digestion and allows for water reabsorption from the kidneys and gastrointestinal tract back to systemic circulation (Rowland, 2007; Watts and Boyle, 2010). However, unlike those studies, we did not observe that the GI tract was empty of food and feces in the mice fed the LF diet suggesting that mice are not limiting food intake. If food intake was restricted we would expect to see genes involved in glycogenolysis and gluconeogenesis (Salter and Watts, 2003; Schoorlemmer and Evered, 2002; Watts and Boyle, 2010), responses that are designed to maintain blood glucose levels (which were significantly different when comparing males and females separately for the SD vs the LFD treatments, Table 1) during fasting or starvation, to be upregulated. While several genes from our consensus results (Supplemental Table 2) were significantly upregulated in LF mice for gluconeogenesis (*PKLR* [hypothalamus], *ADH4, PKLR, HKDC1, ALDOB, PCK1, FBP1* [GI], *PFKFB1* [liver, however this gene inhibits gluconeogenesis], Supplemental Table 2) several were also significantly downregulated in LF mice for gluconeogenesis (*CYP2E1* [liver], *PPP1R3G* [kidney], *ADH4, ENO2,* [GI], Supplemental Table 2). It’s important to note that the two diets differ in the amount of glucose, so understanding if glucose levels and gluconeogenesis are different due to diet or dehydration is not currently possible.

### Diet and the mitochondria

The complex interplay between diet, mitochondrial function, and metabolic homeostasis underscores the dynamic nature of cellular physiology (Dimitriadis et al., 2011). The processing of glucose and fatty acid is controlled via reciprocal regulation between fatty acid oxidation and glucose metabolism, wherein increased glucose consumption suppresses fatty acid oxidation and vice versa (Hue and Taegtmeyer, 2009). The mitochondria play pivotal roles in these regulatory processes, contributing to overall metabolic homeostasis (Caton et al., 2011; Saltiel and Kahn, 2001). When fatty acid oxidation is prioritized, there is an increase in acetyl-CoA and reduced nicotinamide adenine dinucleotide (NADH) levels, which negatively regulates the catalytic activity of PDH through allosteric inhibition and the activation of PDK. During fasting periods, the expression of PDK gene is upregulated by fatty acid-induced peroxisome proliferator– activated receptor (PPAR) signaling (Sugden and Holness, 2006). This regulatory mechanism creates a positive feedback loop that promotes fatty acid oxidation during nutrient scarcity, preserving glucose for essential biosynthetic processes and brain metabolism. Hormones like glucagon-like peptide-1 (GLP-1) and bile acids modulate glucose metabolism and insulin secretion, highlighting the intricate hormonal regulation of metabolic pathways (Jin and Weng, 2016; Kumar et al., 2012; Thomas et al., 2009; van Nierop et al., 2017; Vettorazzi et al., 2016). In this experiment, we saw elevated fatty acid oxidation in the SD, and glucose metabolism in the LFD in mitochondrial genes, however, we also found results contradicting this discussed below.

In the study currently described here there is a robust mitochondrial signal. In the kidney, liver, lung, and hypothalamus several GO terms related to the mitochondria are significantly upregulated for mice fed the low-fat diet. Additionally, we found many overlapping GO terms related to the mitochondria from the independent WGCNA analysis the sex and diet modules in the liver and kidney (inner mitochondrial membrane protein complex [GO:0098800], mitochondrial envelope [GO:0005740], mitochondrial matrix [GO:0005759], mitochondrial inner membrane [GO:0005743], mitochondrial membrane [GO:0031966], mitochondrial translation [GO:0032543], mitochondrion [GO:0005739], and mitochondrial protein-containing complex [GO:0098798], Figure 2) and one GO term in the WLR module for the GI (mitochondrial translation [GO:0032543]. Additionally, *PDK1* and *PDHA1*, convergent genes in cellular energy metabolism (described above) are both significantly differentially expressed in the GI, with *PDHA1* being upregulated in mice fed the LF diet and *PDK1* upregulated in mice fed the SD. Further, 202 genes were significantly downregulated in the LF treatment in the GI and were grouped into the GO term cellular response to insulin stimulus [GO:0032869], as well as nine genes from the consensus GI list (*ADCY2, ATP1A2, HKDC1, KCNMA1, PPP1R3B, PRKCB, RIMS2, RYR2, SNAP25,* Supplemental Table 2) and three genes from the consensus liver list (HNF4A, IGFALS, ITIH2, Supplemental Table 2) related to insulin were downregulated suggesting that the mitochondria are successfully contributing to overall metabolic homeostasis, further explaining the insignificant differences in EE reported in Blumstein et al. (2024).

### Diet, the immune system, and the microbiome

The significant increase in water loss and elevated levels of key electrolytes sensitive to hydration status, including serum Na, osmolality, and Hct, in mice on a LF diet compared to those on a higher-fat diet indicates a distinct difference in water balance between the two diets. As water levels decrease, essential functions, including circulation (Fountain et al., 2023; Georgescu et al., 2017; Santos et al., 2019), temperature regulation (Boonstra, 2004; Canale et al., 2012; Dammhahn et al., 2017; Wingfield et al., 1992), and nutrient transport (Alim et al., 2019; Alonso et al., 2005) is affected. Dehydration-induced stress can activate a stress response and suppress immune system function, impairing the body’s ability to fight off infections and pathogens (Mitchell et al., 2002; Sies, 2015).

Further, diet plays a dual role as both fuel sources and modulators of both the gastrointestinal tract and the microbiome (Henderson et al., 2015; Keeney and Finlay, 2011; Wu et al., 2011). Fatty acids enhance gene expression in innate immune cells (Holzer et al., 2011; Wong et al., 2009; Zeng et al., 2016; Zhao et al., 2007). We observed several genes in our consensus gene lists related immune response upregulated in mice fed a SD (*CNTN2*, *NFASC*, *CD1D*, *CCL11* [lung], *A1BG, C8A, C9, ITIH4, PCDHGB1, ULBP1* [liver], *PTPRZ* [hypothalamus], *PIGR* [kidney], *IL7R* [GI] Supplemental Table 2) and several genes related to inflammation upregulated in mice fed a SD (*ITIH1, CCL11,* [lung], and *ITIH1* [liver], and *ADCY8*, *ASIC4*, *PRKCB* [GI] Supplemental Table 2). However, it’s important to note that inflammation related genes were only identified in three of the five tissues, suggested an isolated responses to altered fat content.

Beyond their inflammatory effects, fatty acids influence bacterial survival and proliferation in the gut (Zeng et al., 2016), partly by disrupting microbial cell membranes (Chen et al., 2011). Consequently, dietary lipid composition can impact gut microbiota colonization and diversity (Jumpertz et al., 2011; Wu et al., 2011), and changes in gut bacteria composition exert ongoing selective pressure on immune modulation by fat (Hooper et al., 2012; Karlsson et al., 2013). The GI, lung, and liver all had significant a WGCNA module with downregulated genes in the LF treatment that grouped into GO terms related to a defense response and or a response to other organisms (regulation of defense response [GO:0031347], response to other organism [GO:0098542], defense response to virus [GO:0051607], response to bacterium [GO:0009617], defense response to symbiont [GO:0140546], Figure 2). An unsuccessful response to infections has been shown to alter behavior in other studies (Cavadini et al., 2007) resulting in the downregulation of clock gene expression and disruption of the circadian clock, leading to a reduction in the amplitude of circadian rhythms (discussed below). Future studies should explore shifts in gut microbiome community structure.

### Diet and circadian rhythm

Circadian rhythmicity of metabolism profoundly influences organ function and systemic homeostasis and in mammals, with the most robust circadian cues being light and food (Pittendrigh 1960; Roenneberg and Merrow 2005). The circadian regulation of physiological processes involves intricate interactions between the central clock in the hypothalamus and peripheral clocks in various organs (Hirota and Fukada, 2004; Xie et al., 2019). While light controls the central clock, peripheral clocks are influenced by rhythmic feeding and fasting cycles, nutrient levels, energy availability, redox levels, and diet composition (Kohsaka et al., 2007; Leone et al., 2015; Tahara et al., 2018; Voigt et al., 2014; Zarrinpar et al., 2014). Metabolic parameters play crucial roles in feedback regulation, affecting core function (Kohsaka et al., 2007; Leone et al., 2015; Zarrinpar et al., 2014). Lipid absorption in the small intestine, regulated by clock genes, displays diurnal variations affected by light and feeding cues (Pan and Hussain, 2009). In our consensus gene sets (Supplemental Table 2) we observed several genes whose primary function delt with digestion and absorption. Interestingly, genes in the consensus GI list related to fat digestion and absorption were upregulated in mice fed the LF diet (*FABP1, PNLIPRP2, ABCG8, FABP2, ABCG5, PLA2G12B, NPC1L1, MTTP, APOA4, APOB,* Supplemental Table 2) while genes in the consensus GI list related to carbohydrate digestion and absorption were upregulated in mice fed the SD (*PRKCB, HKDC1, SLC2A2, ATP1A2, PDHA1* Supplemental Table 2).

The kidney, liver, and GI exhibit rhythmic regulation of blood flow, urine production, and metabolic pathways, modulating processes like detoxification, glucose homeostasis, and lipid metabolism (Solocinski and Gumz, 2015; Zwighaft et al., 2016). Also, gluconeogenesis and glycogen synthesis are under circadian control, involving interactions between core clock components and metabolic regulators (Lamia et al., 2008). Mitochondrial dynamics and function also display circadian rhythmicity, regulated by clock genes (Jacobi et al., 2015; Kohsaka et al., 2014; Magnone et al., 2015) and influenced by diet composition and feeding time (Neufeld-Cohen et al., 2016). Lipids, essential for mitochondrial membrane integrity and energy production, exhibit circadian accumulation patterns (Aviram et al., 2016), synchronized with the molecular clock and feeding cues (Osman et al., 2011).

We found several overlapping GO terms related to circadian rhythm from the independent WGCNA analysis in the kidney, liver, and lung (circadian rhythm [GO:0007623], entrainment of circadian clock by photoperiod [GO:0043153], photoperiodism [GO:0009648], positive regulation of response to stimulus [GO:0048584], and regulation of circadian rhythm [GO:0042752], circadian regulation of gene expression [GO:0032922], circadian behavior [GO:0048512], circadian regulation of translation [GO:0097167], entrainment of circadian clock [GO:0009649], entrainment of circadian clock by photoperiod [GO:0043153], Figure 2). Further, several genes in the consensus GI list related to circadian rhythm downregulated in mice fed the LF diet (*GNAO1, RYR2, GNG3, PRKCB, ADCY2, CACNA1G, GRIN1* Supplemental Table 2). It’s important to note that the core clock genes in the kegg pathway circadian rhythm (hsa04710, Kanehisa et al., 2023) *CLOCK*, *ARNTL* (coded as *BMAL1*), *PER*, *CRY*, *ROR*, and *REV-ERB* were not significant differentially expressed in most tissues, with the exception of *CRY1* and *PER1* in the GI and *ARNTL* and *CRY1* in the lung. Nevertheless, many of these genes were present in significant WGCNA modules and while not significantly differentially expressed they were downregulated in the LF diet compared to the SD. This could suggest these genes may have small changes in gene expression but big effects or other genes that display circadian rhythmicity are driving the expression patterns in the WGCNA analysis (Figure 2).

## Conclusion

We identify several responses employed by the cactus mouse to altered dietary fat content, providing insights into physiological adaptations to increasingly water-stressed plant-based food resources due to climate change. We describe 1) gene expression changes in immune system genes, 2) gene expression changes in circadian regulating genes, and 3) expression changes related to mitochondrial function when comparing cactus mice fed a LF diet to those fed a SD. Our results suggest that a LF diet could limit the capacity of desert animals to tolerate reduced access to free water, as is common in arid environments, as well as pave the way for further exploration and a holistic comprehension of diet’s impact on survival strategies in harsh habitats and the potential implications for long-term population viability amid changing climates.

The interplay between metabolism, diet, the immune response, and circadian rhythms is complex and multifaceted, with implications for overall health and disease. Understanding these interactions sheds light on the intricate regulatory mechanisms that govern physiological processes in a time-dependent manner. There is a push to incorporate -omics with physiology to further understand these processes but currently there are challenges. The analysis discussed is a single contemporary “snapshot” of gene expression. This leaves a mismatch in timescales for contemporary studies as the desired physiological datasets are continuous and contain temporal patterns while gene expression studies that include temporal variation may quickly become overwhelming in cost, scale, and analytical complexity. While we were able to identify genes and GO terms related to circadian rhythm, we don’t yet have data to confirm this result and future studies should explore this. Additionally, grouping genes by function for comparison between organ systems is further complicated due to the hierarchical nature of gene ontology terms. Functional overlap could be obscured by a parent term being assigned to one tissue while a child term is assigned to another. Lastly, we were able to identify gene expression differences in immune system genes for mice on the different diets and a very large change in gene expression in the GI. We hypothesize that the microbiome is playing a role in this response and encourage further studies to characterize differences in the microbiome. Further knowledge of the underpinnings of the physiological phenotypes will be instrumental in discovering more mechanistic links. This will enable a shift from describing correlation to identifying causation of the diverse phenotypes that contribute to phenotypic variation and rapid response to environmental change and combining techniques may allow for a more complete picture of ecological interactions and evolutionary processes.

## Acknowledgments

We thank the members of the MacManes lab for helpful comments and support on previous versions of the manuscript; Adam Stuckert at the University of Huston for lively discussion, valuable insight, and code development; The Animal Resources Office and veterinary care staff at the University of New Hampshire for colony maintenance and care. This work was supported by the National Institute of Health National Institute of General Medical Sciences (R35 GM128843 to M.D.M.).

## Author Contributions

Conceptualization: M.D.M.; Methodology: D.M.B.; Formal analysis: D.M.B., Investigation: D.M.B., Resources: M.D.M.; Writing - original draft: D.M.B.; Writing - review & editing: D.M.B., M.D.M.; Visualization: D.M.B; Supervision: M.D.M.; Project administration: M.D.M.; Funding acquisition: M.D.M.

## Data Availability

Raw reads are available under BioProject ID: PRJNA1048512, SUB14011948 or at <u>http://www.ncbi.nlm.nih.gov/bioproject/1048512</u>. All R scripts used in this project are available through GitHub at: https://github.com/DaniBlumstein/diet_rnaseq.

## Supplemental

**Supplemental Figure 1.**
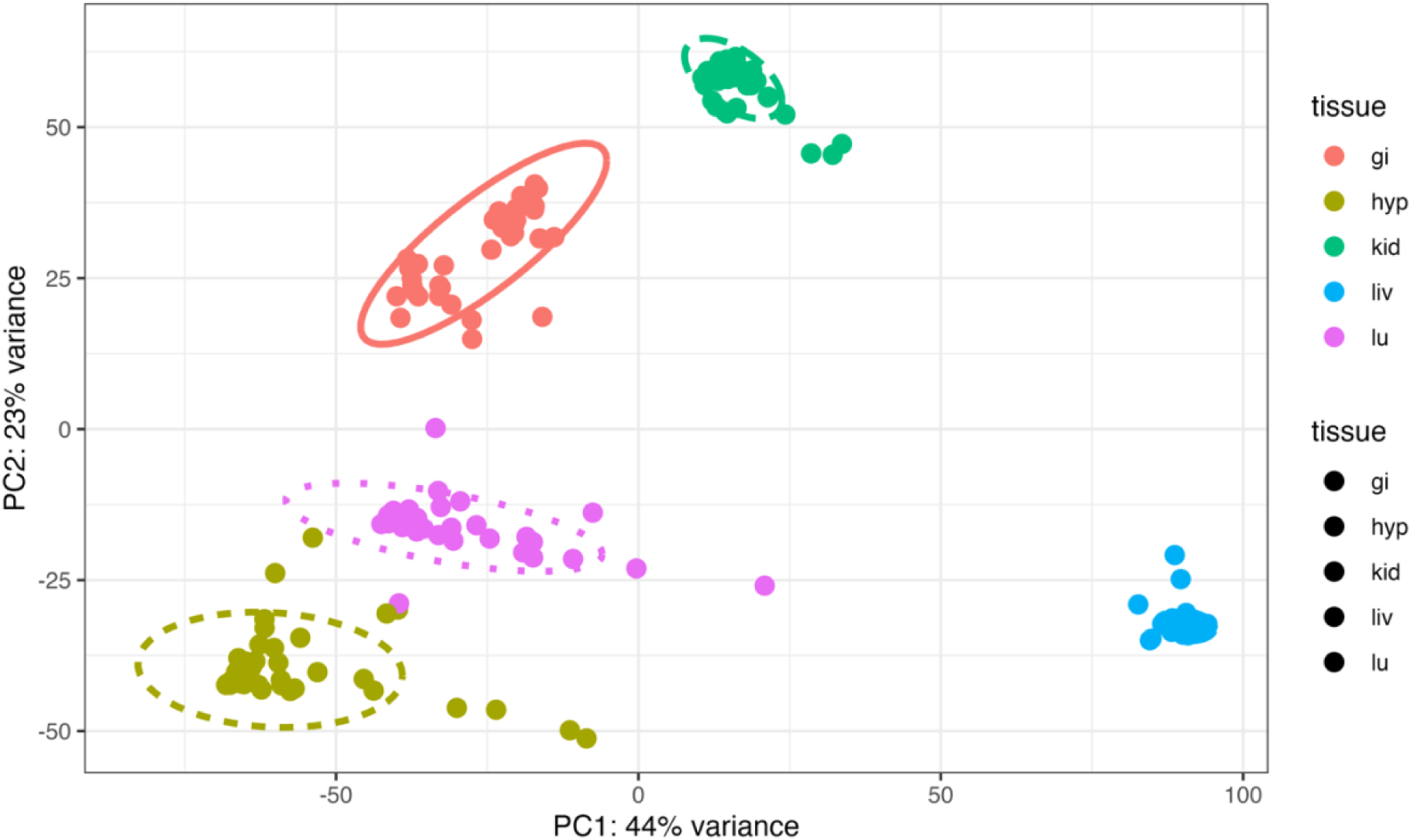
Principal component analysis of gene expression of the lung (lu), liver (liv), gastrointestinal tract (gi), hypothalamus (hyp), and kidney (kid) of *Peromyscus eremicus*. The axes are labelled with the proportion of the data explained by principal components 1 and 2.

Supplemental Table 1

Mapping rates for all samples sequences.

Supplemental Table 2

Genes identified in all three analyses (i.e. significantly differentially expressed genes, assigned to a significant module in WGCNA, and are outliers in CCA) for the lung (lu), liver (liv), hypothalamus (hyp), kidney (kid), and gastrointestinal tract (gi) grouped by the general function of the gene based on GO analysis and KEGG analysis.

